# ADAPT: Analysis of Microbiome Differential Abundance by Pooling Tobit Models

**DOI:** 10.1101/2024.05.14.594186

**Authors:** Mukai Wang, Simon Fontaine, Hui Jiang, Gen Li

## Abstract

Microbiome differential abundance analysis remains a challenging problem despite multiple methods proposed in the literature. The excessive zeros and compositionality of metagenomics data are two main challenges for differential abundance analysis. We propose a novel method called “analysis of differential abundance by pooling Tobit models” (ADAPT) to overcome these two challenges. ADAPT uniquely treats zero counts as left-censored observations to facilitate computation and enhance interpretation. ADAPT also encompasses a theoretically justified way of selecting non-differentially abundant microbiome taxa as a reference for hypothesis testing. We generate synthetic data using independent simulation frameworks to show that ADAPT has more consistent false discovery rate control and higher statistical power than competitors. We use ADAPT to analyze 16S rRNA sequencing of saliva samples and shotgun metagenomics sequencing of plaque samples collected from infants in the COHRA2 study. The results provide novel insights into the association between the oral microbiome and early childhood dental caries.

## 1 Introduction

The microbiome plays an essential role in human health and disease. Extensive research has been conducted into the human microbiome using high-throughput metagenomic sequencing technologies [1–3]. The metagenomics data are count tables that represent each sample’s abundance profiles of microbiome taxa. Differential abundance analysis (DAA) identifies taxa whose abundances differ between conditions. This is one of the fundamental analyses of microbiome data [4]. Many methods have been proposed to tackle statistical challenges in identifying differentially abundant (DA) taxa. However, there is not a universally preferred solution yet [5, 6].

Metagenomics count data have excessive zeros [7, 8]. As illustrated in the toy example in Figure 1a, zeros might reflect the actual absence of taxa in one condition (biological zeros) or indicate rare taxa that the sequencing instrument can not detect (sampling zeros). Some DAA methods impute the zeros with a small positive constant [9–13]. The imputations pave the way for applying standard statistical models to log counts. However, these imputations assume that all zeros are sampling zeros and ignore that library sizes vary among samples, which may lead to inflated false discovery rates [5, 6]. Other methods fit statistical distributions to the counts or count proportions, then draw from the fitted distributions to retrieve smoothed counts and count proportions for downstream analysis [6, 14, 15]. These methods have better control of false discovery rates [6], but their distribution choices lack justification. A third group of methods adopt zeroinflated distributions [16, 17]. Zero-inflated distributions agree with the sparsity patterns in metagenomics data. However, fitting a zero-inflated distribution involves estimating both the probability of true zeros and the distribution parameters for nonzero values. Combining two hypothesis tests for differential abundance analysis reduces power and inflates false discovery rates.

**Fig. 1.**
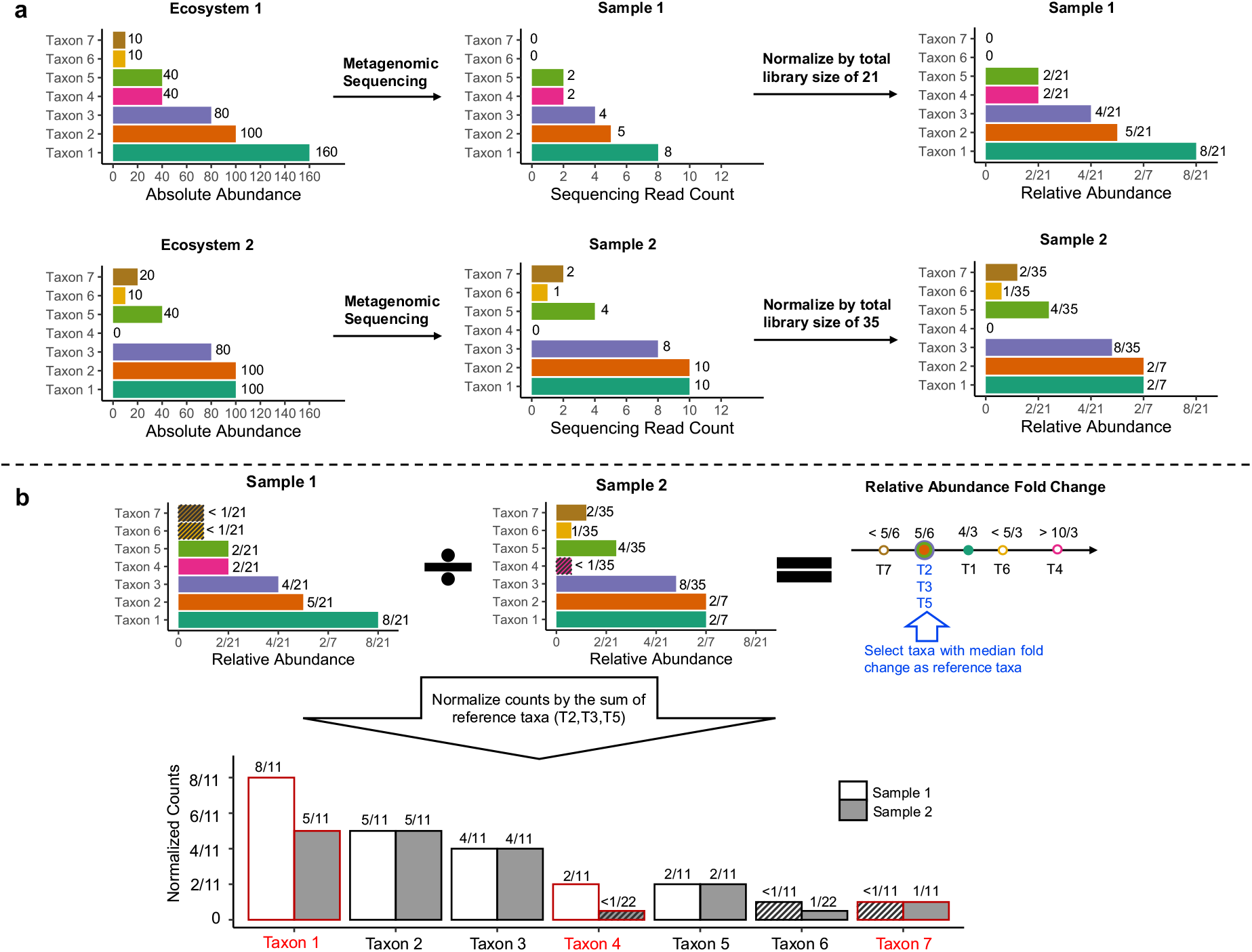
Illustration of ADAPT with a toy example. **(a)** Three microbiome taxa (taxon 1, taxon 4, and taxon 7) are differentially abundant between two ecosystems. Neither the observed counts nor the relative abundances can be directly compared for differential abundance analysis. (**b**) ADAPT treats zero counts as left-censored at the detection limit (one in this case). ADAPT first calculates the fold change of relative abundances. It then selects a subset of taxa (taxon 2, 3, and 5) whose fold changes equal the median as reference taxa. After scaling the counts by the sum of three reference taxa, ADAPT can recover the DA taxa without false positives by comparing the normalized counts.

Metagenomics data are also compositional [18]. As illustrated in Figure 1a, the metagenomics sequencing read counts are not directly comparable between samples because of different sequencing depths. The counts can be interpretable after they are scaled by the library sizes and transformed into relative abundances. However, taxa may have different relative abundances between conditions while their absolute abundances remain stable. The key to DAA is to find an appropriate scaling factor to bridge the gap between relative and absolute abundances. Some methods normalize the counts with centered log-ratio (CLR) transformation [14]. The CLR transformation uses the geometric mean of all taxa counts as the scaling factor. The geometric mean calculation involves DA taxa, which could yield false positives. Other methods derive bias correction factors and have them multiplied with relative abundances or added to CLR transformed counts [10–12, 16]. The estimation procedures of these bias correction factors are derived based on distribution assumptions of absolute abundance fold changes among all the taxa. This strategy is effective when the distribution assumptions are close to the truth. Otherwise, it will lead to high false discovery rates. A third group of methods normalize counts with one or multiple reference taxa [6, 15, 17]. The reference taxa are assumed to be not differentially abundant. This idea is simple and effective [19]. However, existing reference taxa selection procedures rely on calculating log count ratios between taxa pairs, which is computationally expensive and inaccurate given the excessive zero values.

We have developed a new DAA method called “Analysis of Microbiome Differential Abundance by Pooling Tobit Models” (ADAPT). ADAPT has two innovations. First, we treat the zero counts as left-censored at the detection limit of the sequencing instruments. A censored observation unifies different zero mechanisms and accurately reflects the information contained in the observation. Second, we introduce an innovative way of finding non-differentially abundant (non-DA) reference taxa and using reference taxa to identify differentially abundant ones. Under the common assumption that DA taxa are the minority [6], we provide solid theoretical justification that selecting reference taxa is feasible based on all taxa’s relative abundances. We implement these two ideas by incorporating the Tobit model [20] from econometrics and survival analysis. We generate synthetic microbiome count data from independent simulation frameworks and show that ADAPT has better control of false discovery rates and higher power than competitor methods. We also demonstrate our method on the saliva and plaque samples of infants in the COHRA2 [21] study to reveal differentially abundant taxa between kids who developed early childhood dental caries and those who did not.

## 2 Results

### 2.1 ADAPT Workflow

We illustrate the workflow of ADAPT using a toy example (Figure 1). There are seven taxa, and three of them are differentially abundant between two ecosystems. We aim to identify the three DA taxa based on the observed counts in the two metagenomics samples. We denote the counts of undetected taxa 6 and 7 in sample one as left-censored at one (the detection limit, which is assumed to be known). Therefore, their relative abundances are left-censored at 1/21. Similarly, we denote the count of taxon 4 in sample two as left-censored at one and its relative abundance to be left-censored at 1/35. We calculate the relative abundance fold change between two samples for all the taxa. According to the assumption that a minority of taxa are differentially abundant, we can be confident that taxa with median relative abundance fold changes are not differentially abundant (see Methods and Section S1 of Supplementary Materials for theoretical justifications). We choose these taxa as reference taxa. The reference taxa are taxon 2, 3, and 5 in this example. We normalize the individual taxa counts with the sum of reference taxa. By comparing the normalized counts between the two samples, we can correctly identify taxon 1, 4, and 7 as differentially abundant without including false positives.

When analyzing real-life microbiome data with more samples and taxa, we introduce Tobit models for modeling potentially left-censored relative abundances and normalized counts. We pool the effect size estimates and the hypothesis test *p*-values of Tobit models to find reference taxa and identify DA taxa. The detailed procedures of ADAPT are described in Methods.

### 2.2 Performance Evaluation with Simulation Studies

We evaluate the false positive rate control, false discovery rate control, and statistical power of ADAPT with synthetic data. We generate synthetic data using the simulation framework SparseDOSSA [22]. SparseDOSSA framework draws absolute abundances of taxa from zero-inflated log-normal distributions and draws library sizes of all samples from log-normal distributions. The metagenomics sequencing counts are drawn from multinomial distributions based on the simulated library sizes and absolute abundances. The parameters for the simulations are estimated from 16S rRNA sequencing of stool samples in the Human Microbiome Project [23]. We prepare accompanying metadata with a binary covariate and a continuous covariate. The binary covariate represents two contrasting conditions. The zero inflation probabilities and the means of lognormal distribution correlate with this binary covariate for DA taxa. The continuous covariate is a potentially confounding variable. It may be correlated with the metadata’s binary variable of interest and some taxa’s absolute abundances. The details of the simulation setup are described in Methods.

We compare the performance of ADAPT with eight other DAA methods. The competitors are ALDEx2 [14], MaAsLin2 [13], metagenomeSeq [16], ANCOM [9], ZicoSeq [6], DACOMP [15], ANCOMBC [10] and LinDA [12]. These competitors represent a variety of solutions to the excessive zeros and compositionality of metagenomics data (Supplementary Table S1). The proportion of DA taxa, sample sizes, fold changes of absolute abundances, library sizes, and confounding covariates could impact the performance of DAA methods. Therefore, we prepare various scenarios to study their influence. When investigating the influence of one factor, all other factors are fixed. We prepare 500 replicates for each simulation setting and report the average of performance metrics.

We first evaluate the false positive rates (type I errors) of all the DAA methods when there are no differentially abundant taxa between the two conditions (Figure 2a). Because ANCOM does not report raw p-values for DAA, it is excluded from this comparison. When the average library sizes are the same between the two conditions, all the DAA methods can control the FPRs at or below the nominal level of 0.05. When the library sizes are unbalanced, the proportion of sampling zeros and biological zeros differ between conditions. Competitors such as ANCOMBC and LinDA, which replace all the zeros with constant pseudo counts, have the most inflated FPRs. The zero-inflated model by metagenomeSeq also struggles to decipher the zero mechanisms that are confounded by library sizes. DAA methods detect more DA taxa when the sample sizes are larger, which leads to more severely inflated FPRs for some of the competitor methods. ADAPT can maintain FPRs around the nominal level regardless of the unbalanced average library sizes between conditions. This experiment shows that left censoring by ADAPT is more robust than ad hoc zero replacement strategies at handling excessive zeros in metagenomics data.

**Fig. 2.**
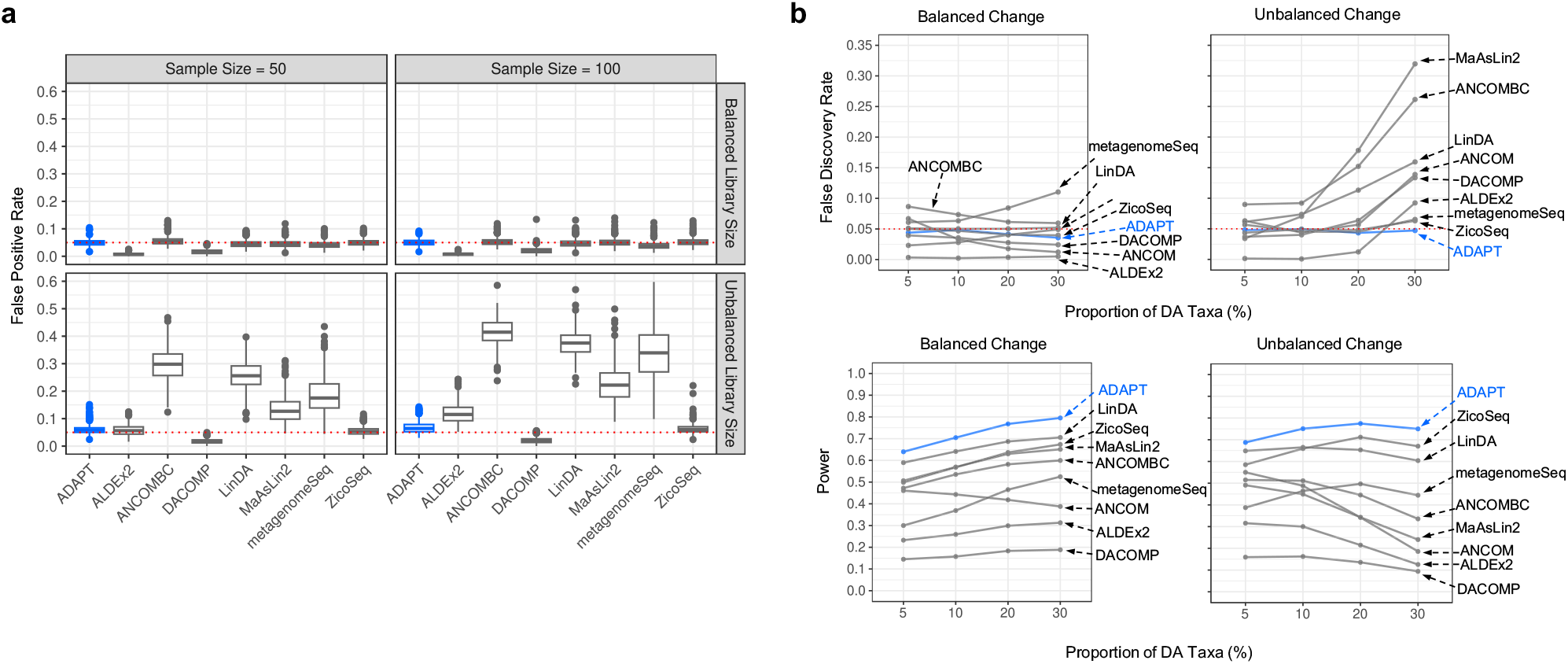
Simulation studies for comparing ADAPT with eight other differential abundance analysis methods. We simulate synthetic metagenomics sequencing count data under two contrasting conditions using the SparseDOSSA framework. The number of samples is the same between the two conditions. We generate 500 replicates for each simulation setting and report the mean of performance metrics. (a) False positive rates (type I errors) of all methods except for ANCOM under simulation settings with no DA taxa. The total number of taxa is 500. The total sample size is 50 or 100. The average library size is the same (balanced) for two conditions at 10^4^ or different (unbalanced) between two conditions (10^4^ for one condition and 10^5^ for the other). (b) False discovery rates and power under simulation settings with different proportions of DA taxa. The sample size is 100. The total number of taxa is 500. The proportion of DA taxa is 5%, 10%, 20%, or 30%. The average library size is 2 × 10^4^ for both conditions. The average absolute abundance fold change of DA taxa is 5. The directions of absolute abundance changes of DA taxa may be balanced or unbalanced.

We then evaluate the false discovery rate control and power of all DAA methods when we experiment with different proportions of DA taxa (Figure 2b). In simulation settings with balanced changes, most DAA methods can control FDR regardless of the proportion of DA taxa. The power of most DAA methods except for ANCOM increases as the proportion of DA taxa increases. Many competitor DAA methods cannot control FDRs in simulation settings with unbalanced changes, especially when DA taxa proportions are high. This is because many normalization strategies by competitor methods assume that the number of taxa whose absolute abundances are enriched is similar to those whose absolute abundances are depleted between two conditions. This assumption is violated when the changes of all DA taxa are in the same direction. ADAPT does not make assumptions about the distributions of absolute abundance fold changes and consistently selects non-DA taxa as reference taxa. The robust selection scheme of reference taxa guarantees false discovery rate control. ADAPT also has the highest average detection power.

Other factors affect the FDR control and power besides the proportion of DA taxa. The detection power of all DAA methods decreases drastically as the sample sizes decrease (Supplementary Figure S1). Several competitor methods, such as metagenomeSeq and ANCOMBC, have inflated FDRs when the sample size is as small as 50. The detection power of all methods increases as the average absolute abundance fold changes of DA taxa increase (Supplementary Figure S2). Increasing the average library sizes of samples boosts the detection power (Supplementary Figure S3). If the absolute abundances are affected by confounding covariates, DAA methods must adjust for the confounders to control FDRs (Supplementary Figure S4). ADAPT has the most consistent control of FDRs and the highest average power across all the simulation scenarios. The computation times of all the DAA methods are mainly determined by the total number of taxa. ADAPT has the best computational efficiency among all the competitors (Supplementary Figure S5). It only takes ADAPT 0.176 seconds to analyze a count table with 1000 taxa and 100 samples. ZicoSeq, which has the best balance of FDR control and power among all competitors, needs 80 seconds.

### 2.3 Oral Microbiome and Early Childhood Dental Caries

Dental caries is the most common chronic disease for US children aged 5 to 17 [24]. Supragingival microbial communities are associated with early childhood dental caries (ECC) [25]. We can use DAA to identify microbiome taxa whose abundances differ between children who developed ECC by five years old and those who did not. We use 16S rRNA sequencing data from saliva samples and shotgun metagenomics sequencing (WGS) data from plaque samples of the Center for Oral Health Research in Appalachia 2 (COHRA2) cohort [26]. There are 161 saliva samples collected at 12 months old. None of the 161 children had dental caries during sample collection. Among these children, 84 later developed ECC, and 77 did not. There are 30 plaque samples collected between 36 and 60 months old. Half of the 30 samples were collected from children with dental caries, and the other half were caries-free. The plaque samples of the cases were collected at the onset of ECC. We remove taxa with prevalence lower than 5%, leading to 155 amplicon sequence variants (ASVs) in the count table of the saliva samples and 590 taxa in the count table of the plaque samples. We apply all the nine methods compared in the simulation studies and carry out DAA for the saliva samples and plaque samples separately. We apply the Benjamini-Hochberg correction to the raw p-values for all the DAA methods and use 0.05 as the cutoff level for DA taxa identification.

Among the 155 ASVs in the saliva samples, 38 are identified as differentially abundant by at least one DAA method (Figure 3a). ADAPT identifies 27 differentially abundant ASVs. Several ASVs discovered by ADAPT were mentioned in multiple previous studies according to a recent review [27], including *Haemophilus parainfluenzae, Fusobacterium periodonticum, Prevotella histicola, Veillonella parvula, Lachnoanaerobaculum umeaense* and *Porphyromonas pasteri*. Among these six species, *H. parainfluenzae, F. periodonticum, L. umeaense*, and *P. pasteri* are enriched in children free of dental caries. *P. histicola* and *V. parvula* are enriched in children who later developed dental caries. These trends align with findings in previous literature as well.

**Fig. 3.**
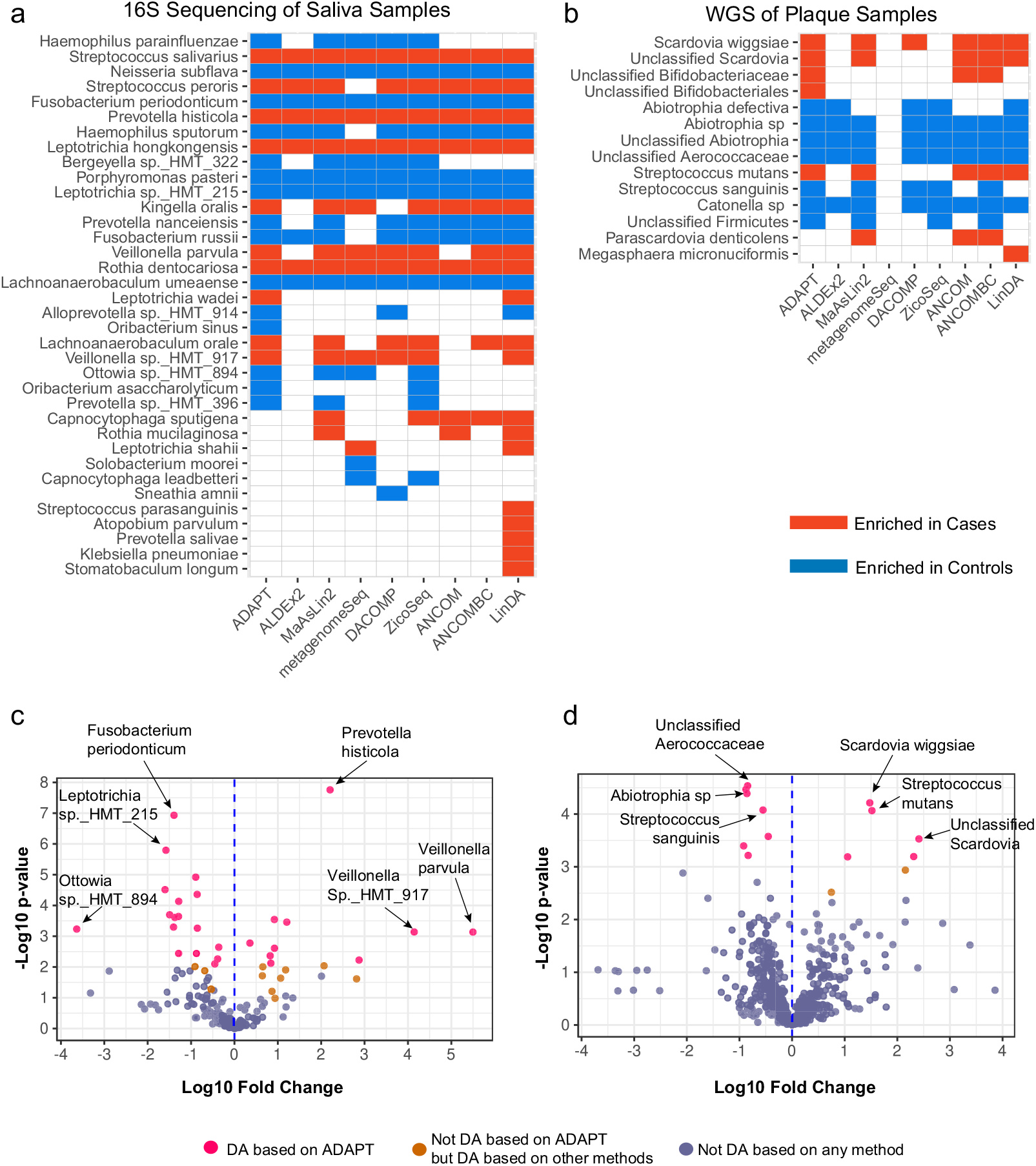
Microbiome differential abundance analysis between children who developed early childhood dental caries and those who did not. (a) 38 out of 155 total amplicon sequence variants in the saliva samples collected at 12 months old are differentially abundant based on at least one method. ADAPT detects 27 DA ASVs. (b) 14 out of 590 taxa in the plaque samples collected between 36 and 60 months old are differentially abundant based on at least one method. ADAPT detects 12 DA taxa. (c) Volcano plot for DAA of saliva samples (d) Volcano plot for DAA of plaque samples

Among the 590 taxa in the plaque samples, 14 are identified as differentially abundant by at least one DAA method (Figure 3b). ADAPT identifies 12 DA taxa. The discoveries of ADAPT include *Scardovia wiggsiae, Streptococcus mutans*, and *Streptococcus sanguinis*, which were mentioned in previous reviews [27]. According to ADAPT, *S. sanguinis* is enriched in controls. *S. mutans* and *S. wiggsiae* are enriched in cases. These trends agree with previous findings as well. The DA taxa in the plaque samples collected after 36 months old differ from the DA taxa in the pre-incident saliva samples collected at 12 months old. This phenomenon echoes the idea that microbiome species associated with dental caries vary with age [26, 27].

ADAPT estimates the absolute abundance fold changes besides identifying DA taxa. Most taxa’s estimated log10 fold changes are between -2 and 2 for both the saliva and the plaque samples (Figures 3c and 3d). For some rare taxa that only exist in samples from one condition, the absolute values of their estimated effect sizes are much larger than the others. For example, *Veillonella parvula* has the largest estimated log10 fold change of 5.50 among all the ASVs in the saliva samples. This species was detected in 10 of the 161 saliva samples, and all these ten samples were from children who eventually developed dental caries. The count proportions of *Veillonella parvula* in these ten samples range from 6 × 10^−4^ to 0.06. Still, complete separation does not necessarily indicate DA taxa. For example, *Fusobacterium naviforme* is detected only in three of the 30 plaque samples, all from children without dental caries. The estimated log10 fold change is -3.687, the lowest among all the taxa. However, the corresponding *p*-value is 0.09, so *Fusobacterium naviforme* is not considered differentially abundant. The count proportions of *Fusobacterium naviforme* among the three samples are only 5.8 × 10^−6^, 1.0 × 10^−6^, and 3.1 × 10^−7^. The computational heuristics in ADAPT prevent infinite log abundance fold change estimates and enable valid statistical tests when complete separation occurs (Section S2 of Supplementary Materials). Analyses of real-life data show that ADAPT is robust and numerically stable for evaluating differential abundance patterns of rare taxa.

## 3 Discussion

The excessive zeros in metagenomics data come from multiple sources, and it is challenging to classify and preprocess them for differential abundance analysis [7, 8, 28]. Left censoring can circumvent any controversial zero preprocessing steps. This idea has only seen limited use in another work that compares relative abundances between conditions [29]. ADAPT is the first method to demonstrate the ingenuity of censoring when comparing absolute abundances. The simulation studies and real data analyses prove that censoring can control false discovery rates and maintain competitive detection power. We choose to censor all the observed zero counts at one (the smallest nonzero value) for all the simulations and real data analyses. This proxy is a natural choice, given that metagenomics counts are discrete. Nevertheless, other choices are worth exploring. For example, we may censor the zeros at the smallest positive value in the count table if the smallest positive value does not equal one. We may also customize sample-specific proxies given the library size and the smallest positive count in each sample. The prevalence of each taxon is another factor worth considering. The assumption that differentially abundant taxa are the minority is sufficient for our reference taxa selection scheme to be admissible. Nevertheless, our selection of reference taxa can still hold when more than half of all the taxa are differentially abundant as long as the directions of changes are balanced (Supplementary Figure S6). In all simulations and real data analyses, reference taxa account for half or one-fourth of all taxa (Supplementary Figure S7). We notice that the selected reference taxa may contain several differentially abundant ones. The criteria for a qualified reference taxa set is that the hypothesis test *p*-values form a uniform or left-skewed distribution (see Methods for details). Because the decision is based on the distribution of *p*-values instead of thresholds for individual *p*-values, a handful of DA taxa is expected to remain in the reference taxa set. A sizable number of taxa are selected as reference, so minor contamination of DA taxa does not affect the performance of ADAPT. Adding up the counts of multiple reference taxa ensures nonzero normalizing when calculating count ratios. It also decreases the variance of the normalizing factor in comparison to using a single reference taxon, leading to increased power.

The Tobit model is similar to the accelerated failure time model in survival analysis except that it models left-censored data. It is a parametric censored quantile regression. The simulation studies and real data analyses demonstrate microbiome differential abundance between two conditions, but ADAPT can also handle continuous conditions and adjust for multiple covariates. In future work, we will accommodate multigroup comparisons and include random effects for longitudinal study designs. Nonparametric censored quantile regressions [30, 31] are viable alternatives to the Tobit model. They could be more robust than the Tobit model when the distribution of log count ratios is very different from a normal distribution or for small sample sizes. Potential downsides of nonparametric models include computational complexity and lower power. The successful implementation of ADAPT offers new perspectives on handling excessive zeros and compositional data for microbiome differential abundance analysis. It is a valuable addition to the field of ecological data analysis.

## 4 Methods

### 4.1 Mathematical Properties of Relative Abundance

The intuition of ADAPT is supported by mathematical properties of relative abundance. We first present four propositions about relative abundance, which set the stage for deriving ADAPT analysis procedures. The proofs for these four propositions are in Section S1 of Supplementary Materials.

Let 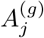 and 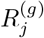 represent the absolute abundance and relative abundance of taxon *j* (*j* = 1, 2, …, *P*) in condition *g* (*g* = 1, 2). The goal of DAA is to decide if 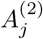 is different from 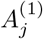 for each taxon *j*. The absolute abundances cannot be directly measured but the relative abundances can be estimated from observed sequencing counts. According to definition, 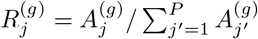.

#### Proposition 1 (Reference taxa)

*Let* T_0_ ⊆ T = {1, 2, · · ·, *P* } *be a set of non-DA taxa. Then, the relationship between the relative abundance fold changes and the absolute abundance fold changes of any taxon j (j* = 1, 2, · · ·, *P ) satisfies*

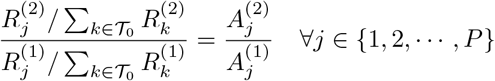

We can calculate the absolute abundance fold change for any taxa from their relative abundances if we find a subset of non-DA taxa as reference taxa. The calculation involves the relative abundance ratio between the taxon of interest and the sum of reference taxa.

#### Proposition 2 (Null Case)

*If none of the taxa* T = {1, 2, · · ·, *P* } *are differentially abundant, then the relative abundances of all the taxa remain the same across conditions. On the other hand, if the relative abundances of all the taxa remain the same between conditions, then the absolute abundance fold changes of all the taxa are the same. Namely*,

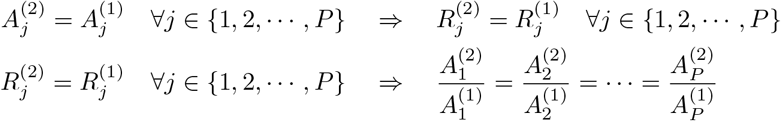

Suppose the relative abundances of all the taxa remain the same between conditions. In that case, all taxa’s absolute abundance fold changes may equal a constant other than one. However, most taxa are assumed to be non-DA [6, 32]. Based on this assumption, we can decide that there are no DA taxa if no relative abundances change between conditions for any taxa.

#### Proposition 3 (Order preservation for abundance fold changes)

*Between any two taxa j and k (*1 ≤ *j < k* ≤ *P ), their order of relative abundance fold changes is the same as their order of absolute abundance fold changes. Namely*

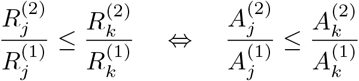

Relative abundance fold change is different from absolute abundance fold change. Still, relative abundance fold change is ranked the same among all taxa as absolute abundance fold change.

#### Proposition 4 (Relative abundance fold change of non-DA taxa)

*Under the assumption that a minority of taxa are DA, the relative abundance fold change of a non-DA taxon j (j* = 1, 2, · · ·, *P ) equals the median of the relative abundance fold changes of all taxa. Namely*

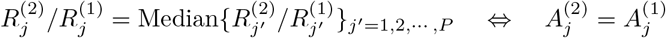

The relative abundances of non-DA taxa differ between two conditions when DA taxa exist. Nevertheless, we can rank the relative abundance fold changes and select taxa with median fold changes as reference taxa.

### 4.2 ADAPT Procedures

ADAPT consists of three main procedures. The first step estimates the relative abundance fold changes of all the taxa with Tobit models and decides if any DA taxa exist. The second step selects a subset of non-DA taxa as reference taxa. This step is needed if the first step confirms the existence of at least one DA taxa. The third step identifies DA taxa by fitting Tobit models to the log count ratios between each taxon and the reference taxa. Supplementary Figure S8 provides a flowchart of ADAPT procedures.

#### 4.2.1 Relative Abundance Fold Change Estimation with Tobit Models

The metagenomics count table *Y* has *N* samples and *P* taxa. The count of taxon *j* (*j* = 1, 2, · · ·, *P* ) in sample *i* (*i* = 1, 2, · · ·, *N* ) is denoted as *y*_*ij*_. Each sample *i* has its vector of covariates *x*_*i*_ including the intercept. The main variable of interest *x*_*i*1_ is binary for DAA between two conditions. If *y*_*ij*_ is zero, we represent it as being left-censored at a positive value *d*

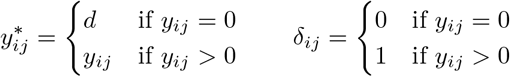

The default value of *d* is one because metagenomic sequencing counts are integers and the detection limit of the sequencing instrument is one. The relative abundance of taxon *j* in sample *i* is denoted 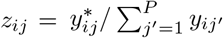. We fit a Tobit model [20] to the log relative abundances {log *z* } of each taxon *j* ∈ {1, 2, · · ·, *P* } by calculating the maximum likelihood estimate of

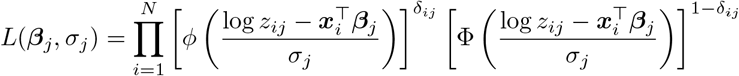

where *Φ*(·) and Φ(·) represent the probability density function and cumulative distribution of the standard normal distribution. 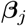 includes the effect sizes of all the covariates. *σ* is the scale parameter that accounts for the variance of log relative abundances. To guarantee the numerical stability of model fitting for rare taxa, we estimate the MLE of the Firth penalized likelihood [33]. Section S2 of Supplementary Materials describes the details of computational heuristics.

We report the effect size estimate 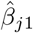 which represents the log relative abundance fold change of taxon *j* between conditions. We also carry out likelihood ratio test *H*_0_ : *β*_*j*1_ = 0 against *H*_1_ : *β*_*j*1_ ≠ 0 and report the *p*-value. We pool the *p*-values of all the Tobit models to find DA taxa in the following steps.

#### 4.2.2. Reference Taxa Selection

If there are no DA taxa, the *p*-values of hypothesis tests for relative abundance fold changes in the first step {*w*_*j*_}_*j*=1,2,···,*P*_ should display a uniform distribution according to Proposition 2. We fit a beta-uniform mixture [34]

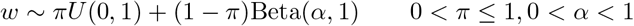

and apply likelihood ratio tests for *H*_0_ : *π* = 1 against *H*_1_ : *π <* 1. If *H*_0_ cannot be rejected, the distribution of {*w*_*j*_}_*j*=1,2,···,*P*_ follows a uniform or left-skewed distribution, indicating that there are no differentially abundant taxa.

If *H*_0_ is rejected, the distribution of p-values is right-skewed and there are DA taxa. We must search for a subset of reference taxa before identifying DA taxa. According to Proposition 3 and 4, a taxon *j* is likely non-DA if its relative abundance fold change estimate 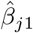 is close to median 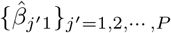 Therefore, we select a subset 𝒯 ^′^ with half of all the taxa whose relative abundance fold change estimates are closest to the median.

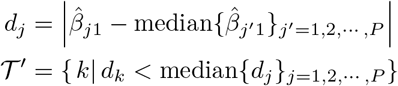

We verify if there are any DA taxa in 𝒯 ^′^ in a way similar to the first step. For each taxon *k* ∈ 𝒯 ^′^, we fit Tobit models to the count proportion within this subset 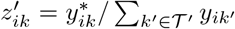. We then check if the distribution of *p*-values from these Tobit models is still right-skewed. If the *p*-value distribution is uniform or left-skewed, 𝒯 ^′^ is a qualified set of reference taxa. Otherwise, we repeat this second step until we obtain a subset of non-DA taxa 𝒯_0_ as reference taxa.

#### 4.2.3 Identification of Differentially Abundant Taxa

Once we identify a subset of non-DA taxa as the reference set, we calculate the count ratio between each taxon *j* and the summed counts of the reference taxa 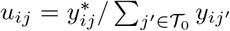 ′ for all samples *i* = 1, 2, · · ·, *N* . We fit a Tobit model to {log *u*_*ij*_}_*i*=1,2,···,*N*_ by calculating the maximum likelihood estimate of

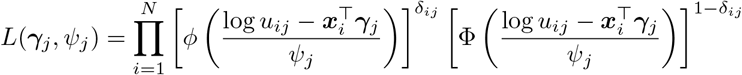

The effect size estimate 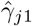 represents the log fold change of absolute abundance for taxon *j* according to Proposition 1. The hypothesis test *H*_0_ : *γ*_*j*1_ = 0 against *H*_1_ : *γ*_*j*1_ ≠ 0 indicates whether taxon *j* is differentially abundant. We apply multiple testing corrections to all the *p*-values to control false discovery rates and call a taxon differentially abundant if its adjusted *p*-value is below a certain level (e.g., 0.05). The simulation studies indicate that the Benjamini-Hochberg method is suitable for multiple corrections.

### 4.3 Simulation Framework

#### 4.3.1 Metadata Generation

The metadata has two variables *X* and *C*. Variable *X*_*i*_(*i* = 1, 2, · · ·, *N* ) is a binary variable representing two contrasting conditions. It is the variable of interest. Variable *C*_*i*_ is a potentially confounding continuous covariate. For each sample *i, X*_*i*_ and *C*_*i*_ are generated in the following way

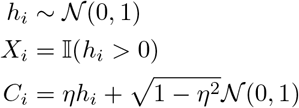

where *η* is the correlation parameter that controls the severity of confounding.

#### 4.3.2 Count Table Generation

SparseDOSSA [22] generates the count table *Y* based on the metadata. The simulation scheme of SparseDOSSA first generates the absolute abundances of each taxon from a zero-inflated log-normal distribution. For taxon *j*(*j* = 1, 2, · · ·, *P* ) in sample *i*(*i* = 1, 2, · · ·, *N* ),

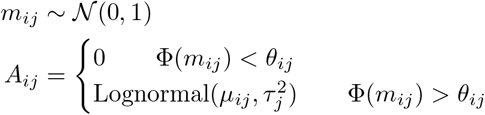

where *m*_*ij*_ is a latent variable for deciding if *A*_*ij*_ *>* 0. The zero inflation probability *θ*_*ij*_ and the log-normal distribution parameter *µ*_*ij*_ depend on *X*_*i*_ and *C*_*i*_

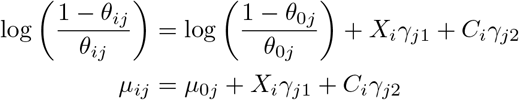

Taxon *j* is differentially abundant if *γ*_*j*1_ ≠ 0. Its abundance is correlated with the potential confounder if *γ*_*j*2_ ≠ 0. The parameters 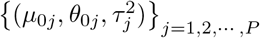 are drawn from the pre-trained template in the SparseDOSSA package. The pre-trained template was calculated based on 16S rRNA sequencing of stool samples in the Human Microbiome Project [1, 23]. There are 332 sets of zero-inflated log-normal distribution parameters in the pre-trained template. DAA performance is indistinguishable among different methods if the simulated count table contains too many rare taxa. Therefore, we only draw (with replacement) from 54 sets of parameters whose zero inflation probabilities are below 50% to set up absolute abundance distributions for all taxa. The relative abundances can be derived based on the generated absolute abundances 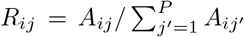. We draw the library sizes from another log-normal distribution and generate the taxon counts from a multinomial distribution

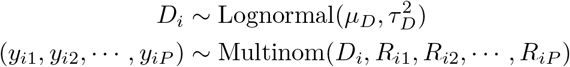

### 4.4 Real Data Preprocessing

The raw sequencing data can be downloaded from NCBI. For the 16S rRNA sequencing data, we filter and trim the reads with low quality. After that, we use the DADA2 [35] package to denoise the reads and assign taxonomy to them based on the HOMD database [36]. For the shotgun metagenomics sequencing data, we use the SqueezeMeta [37] pipeline to carry out genome assembly and taxonomy assignment.

## Supporting information

Source data for Figure 3

## 5 Data Availability

The 16S rRNA sequencing data and shotgun metagenomics sequencing data of the COHRA2 study [26] are available under project number PRJNA752888 (https://www.ncbi.nlm.nih.gov/bioproject/PRJNA752888). The metadata can be requested from the dbGaP database under study accession number phs001591.v1.p1. The preprocessed count tables and de-identified metadata for the metagenomics data in the early childhood dental caries study are available through the ADAPT R package (https://github.com/mkbwang/ADAPT).

## 6 Code Availability

ADAPT is available as an R package on GitHub (https://github.com/mkbwang/ADAPT). The codes for simulation studies, sequencing data preprocessing, and real data differential abundance analysis are also available on GitHub (https://github.com/mkbwang/ADAPT_example).

## 7 Acknowledgements

The authors thank Betsy Foxman and Freida Blostein for providing the metagenomics data and offering guidance on data preprocessing. This research was supported in part through computational resources and services provided by Advanced Research Computing at the University of Michigan, Ann Arbor. M.W. and G.L. were supported by funding from the National Institute of Health under grant number R03DE027773.

## 8 Author Contributions

M.W. and G.L. contributed equally to the methodology development. M.W. was responsible for the method implementation, simulation studies, real data analysis, and manuscript writing. S.F., H.J., and G.L. contributed to the editing of the manuscript.

