## Supplementary Figures, Table and Notes for "ADAPT: Analysis of Microbiome Differential Abundance by Pooling Tobit Models"

### Supplementary Materials for "ADAPT: Analysis of Microbiome Differential Abundance by Pooling Tobit Models"

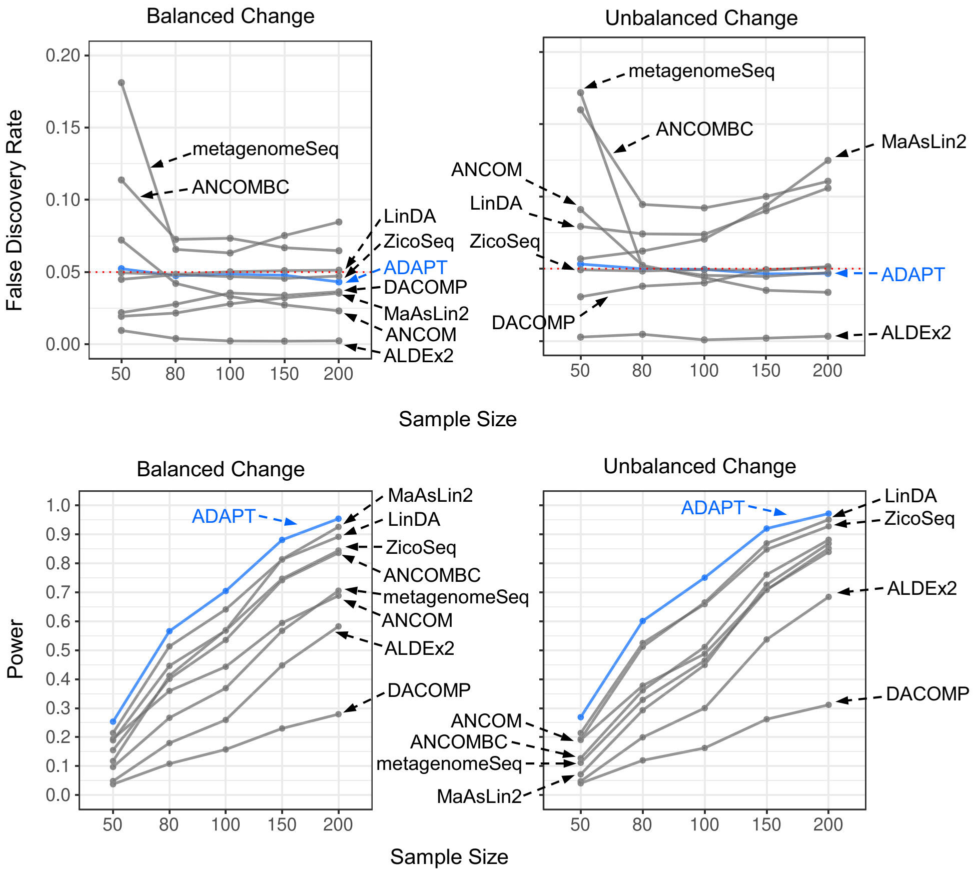


**Fig. S1**: **Simulation studies with different total sample sizes.** The sample size is 50, 80, 100, 150, or 200. The total number of taxa is 500. The proportion of DA taxa is 10%. The average fold change is 5. The average library size is $2 \times 10^4$ for both conditions. The directions of absolute abundance changes of DA taxa may be balanced or unbalanced.

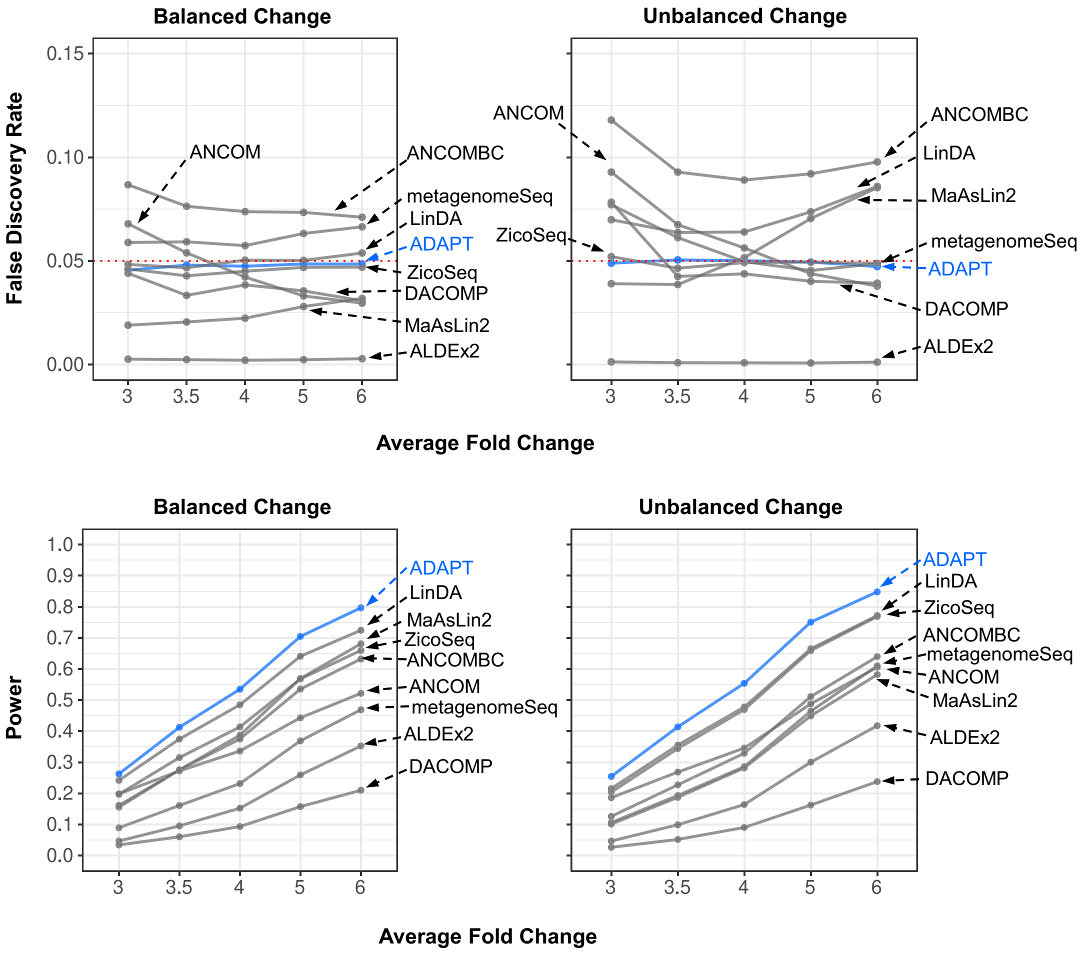


**Fig. S2**: **Simulation studies with different average fold changes of DA taxa.** The sample size is 100. The total number of taxa is 500. The proportion of DA taxa is 10%. The average fold changes are 3, 3.5, 4, 5, or 6. The average library size is $2 \times 10^4$ for both conditions. The directions of absolute abundance changes of DA taxa may be balanced or unbalanced.

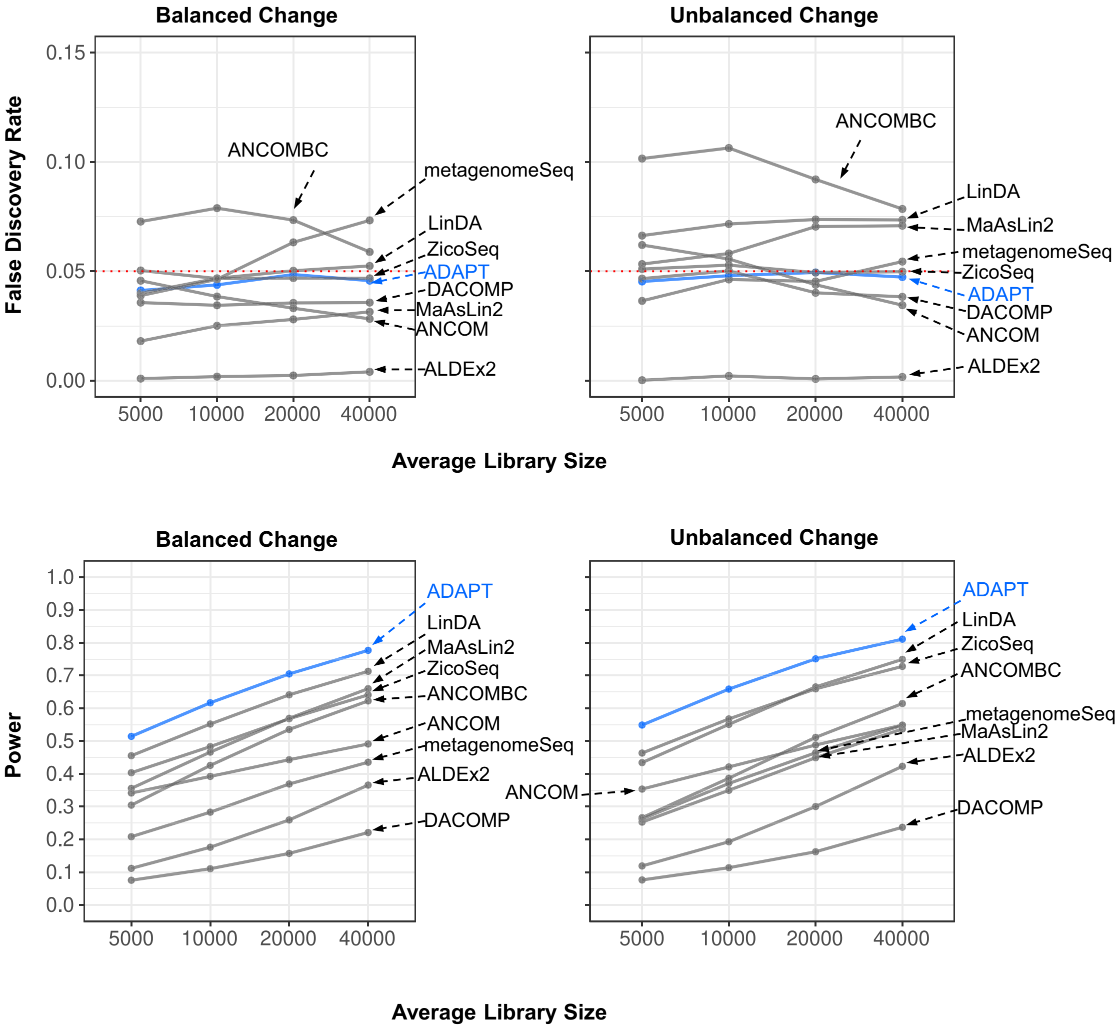


**Fig. S3**: **Simulation studies with different average library sizes.** The sample size is 100. The total number of taxa is 500. The proportion of DA taxa is 10%. The average fold change is 5. The average library size is $5 \times 10^3$, $10^4$, $2 \times 10^4$, and $4 \times 10^4$ for both conditions. The directions of absolute abundance changes of DA taxa may be balanced or unbalanced.

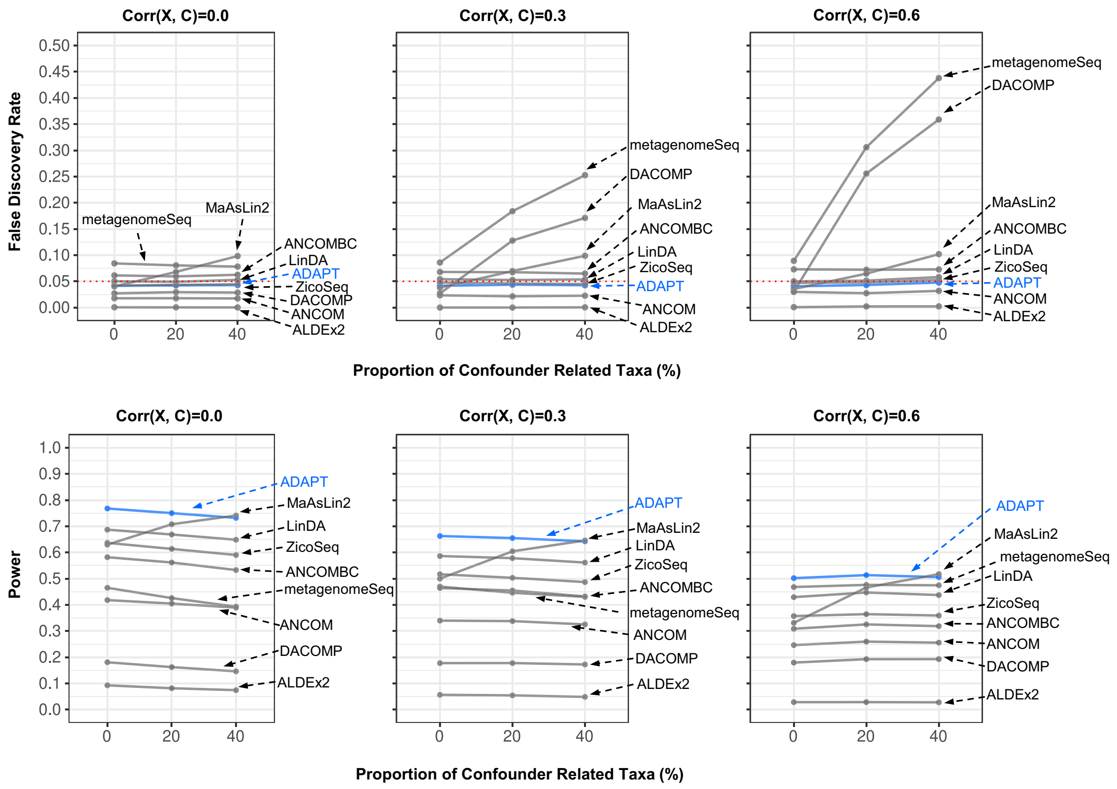


**Fig. S4**: **Simulation studies with potential confounders.** The sample size is 100. The total number of taxa is 500. The proportion of DA taxa is 20%. The average library size is the same for both conditions at $2 \times 10^4$. The average absolute abundance fold change of DA taxa is 5. The proportion of taxa whose abundances correlate with the confounding variable is 0%, 20%, or 40%. The correlation between the binary variable and the continuous confounding variable is 0, 0.3, or 0.6. DACOMP and metagenomeSeq are the only two DAA methods that can not adjust for covariates.

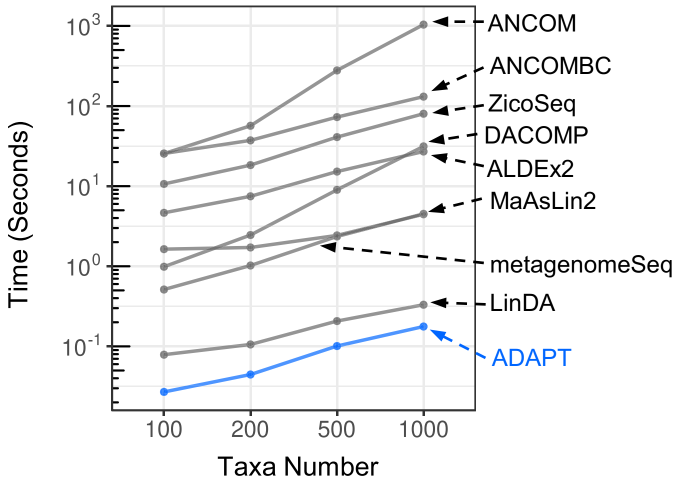


**Fig. S5**: **Computation time (seconds).** All simulations for time measurement have 100 samples and an average library size of $2 \times 10^4$. The total number of taxa is 100, 200, 500, or 1000. 10% of all the taxa are DA. We generate 500 replicates for each simulation setting and report all DAA methods' mean computation time. Each method is allocated four cores (Intel Xeon Gold 6154) and 16GB of memory.

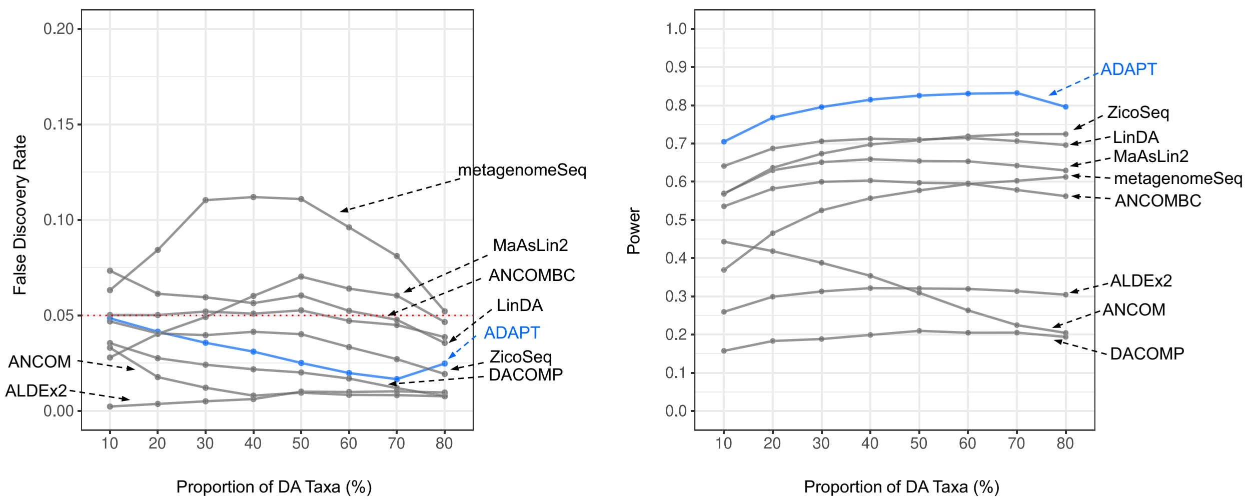


**Fig. S6**: If the assumption that differentially abundant taxa are the minority is violated, ADAPT can still control false discovery rates when the direction of changes is balanced.

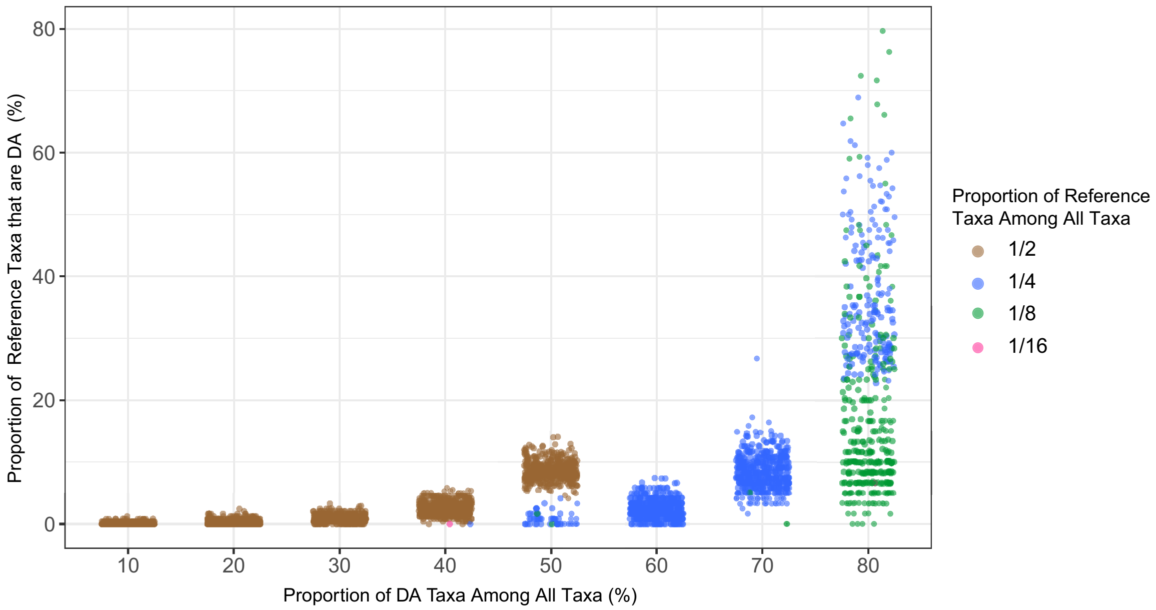


**Fig. S7**: We prepare 500 replicates for each simulation setting with different proportions of DA taxa among all the taxa. The total number of taxa is 500. The directions of change of DA taxa are balanced. ADAPT selects half of all taxa as reference taxa when the proportion of DA taxa among all taxa is below 50%. When more than half of all taxa are differentially abundant, ADAPT will select fewer reference taxa to guarantee low contamination of DA taxa in the reference set. The reference taxa selection scheme stays robust when the proportion of DA taxa is as high as 70%.

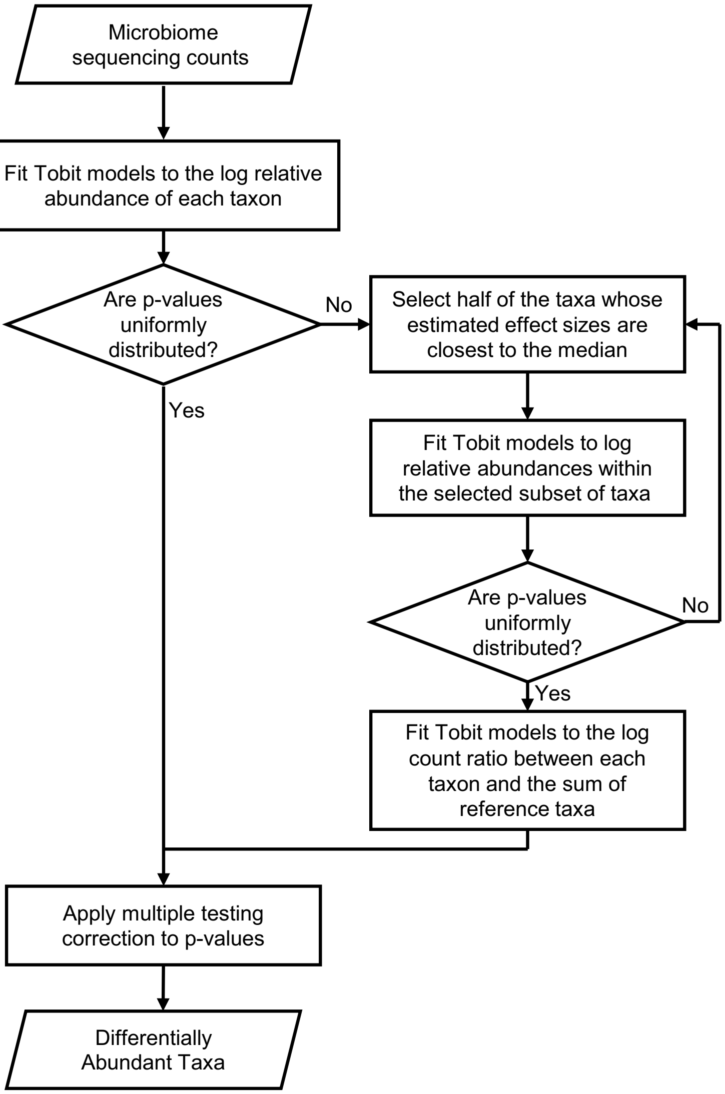


**Fig. S8**: **Analysis procedures of ADAPT.** The first step of ADAPT is to fit Tobit models to the log relative abundances of all the taxa. The second step is to find a subset of non-DA taxa as reference taxa. The third step is to find differentially abundant taxa by fitting Tobit models to log count ratios between individual taxa and the summed counts of reference taxa.

| Method | Compositionality Handling | Zero Handling | Statistical Model |
|---|---|---|---|
| ADAPT | Reference taxa | Left censoring | Tobit model |
| ALDEx2 [1] | Centered log-ratio | Fit Dirichlet distribution and draw count proportions | Wilcoxon test/GLM |
| MaAsLin2 [2] | Total sum scaling | Impute with pseudo counts | Log-linear model |
| metagenomeSeq [3] | Wrench (compositionality correction factor for relative abundances) | Zero-inflated model | Zero-inflated log-normal |
| ANCOM [4] | Count ratios of all taxa pairs | Impute with pseudo counts | Log-linear model |
| DACOMP [5] | Reference taxa | Fit hypergeometric distribution and draw counts | Wilcoxon test |
| ZicoSeq [6] | Reference taxa | Fit beta mixture distribution and draw count proportions | Linear model for power transformed counts |
| ANCOMBC [7, 8] | Bias correction factor for CLR transformed counts | Impute with pseudo counts | Linear model for CLR transformed counts |
| LinDA [9] | Bias correction factor for CLR transformed counts | Impute with pseudo counts | Linear model for CLR transformed counts |

**Table S1**: The solutions of nine differential abundance analysis methods for compositional metagenomics data with excessive zero counts

### S1 Proofs for Propositions

This section contains the proofs for the four propositions about relative abundance.

*Proof for Proposition 1.* We know that $R_j^{(g)} = A_j^{(g)} / \sum_{j'=1}^{P} A_{j'}^{(g)}$ and $\sum_{k\in\mathcal{T}_0} R_k^{(g)} = \sum_{k\in\mathcal{T}_0} A_k^{(g)} / \sum_{j'=1}^{P} A_{j'}^{(g)}$, therefore

$$\frac{R_j^{(g)}}{\sum_{k\in\mathcal{T}_0} R_k^{(g)}} = \frac{A_j^{(g)}}{\sum_{k\in\mathcal{T}_0} A_k^{(g)}}$$

Furthermore

$$\frac{R_j^{(2)} / \sum_{k\in\mathcal{T}_0} R_k^{(2)}}{R_j^{(1)} / \sum_{k\in\mathcal{T}_0} R_k^{(1)}} = \frac{A_j^{(2)} / \sum_{k\in\mathcal{T}_0} A_k^{(2)}}{A_j^{(1)} / \sum_{k\in\mathcal{T}_0} A_k^{(1)}}$$

Because all the taxa in $\mathcal{T}_0$ are non-DA, $\sum_{k\in\mathcal{T}_0} A_k^{(2)} = \sum_{k\in\mathcal{T}_0} A_k^{(1)}$, therefore

$$\frac{R_j^{(2)} / \sum_{k\in\mathcal{T}_0} R_k^{(2)}}{R_j^{(1)} / \sum_{k\in\mathcal{T}_0} R_k^{(1)}} = \frac{A_j^{(2)}}{A_j^{(1)}}$$

□

*Proof for Proposition 2.* If $A_j^{(2)} = A_j^{(1)} \quad \forall j \in \{1, 2, \cdots, P\}$, then the total microbial loads are the same between two conditions as well, namely $\sum_{j'=1}^{P} A_{j'}^{(2)} = \sum_{j'=1}^{P} A_{j'}^{(1)}$. We can deduce that

$$R_j^{(2)} = \frac{A_j^{(2)}}{\sum_{j'=1}^{P} A_{j'}^{(2)}} = \frac{A_j^{(1)}}{\sum_{j'=1}^{P} A_{j'}^{(1)}} = R_j^{(1)}$$

On the other hand, if we know that the relative abundance $R_j^{(2)} = R_j^{(1)} \quad \forall j \in \{1, 2, \cdots, P\}$, we can get

$$\frac{A_j^{(2)}}{\sum_{j'=1}^{P} A_{j'}^{(2)}} = \frac{A_j^{(1)}}{\sum_{j'=1}^{P} A_{j'}^{(1)}} \quad \Rightarrow \quad \frac{A_j^{(2)}}{A_j^{(1)}} = \frac{\sum_{j'=1}^{P} A_{j'}^{(2)}}{\sum_{j'=1}^{P} A_{j'}^{(1)}} \quad \forall j \in \{1, 2, \cdots, P\}$$

This indicates that $\frac{A_1^{(2)}}{A_1^{(1)}} = \frac{A_2^{(2)}}{A_2^{(1)}} = \cdots = \frac{A_P^{(2)}}{A_P^{(1)}}$ □

*Proof for Proposition 3.* According to the definition of relative abundance,

$$\frac{R_j^{(2)}}{R_j^{(1)}} = \frac{A_j^{(2)} / \sum_{j'=1}^{P} A_{j'}^{(2)}}{A_j^{(1)} / \sum_{j'=1}^{P} A_{j'}^{(1)}} \qquad \frac{R_k^{(2)}}{R_k^{(1)}} = \frac{A_k^{(2)} / \sum_{j'=1}^{P} A_{j'}^{(2)}}{A_k^{(1)} / \sum_{j'=1}^{P} A_{j'}^{(1)}}$$

Therefore

$$\frac{R_j^{(2)}}{R_j^{(1)}} < \frac{R_k^{(2)}}{R_k^{(1)}} \quad \Leftrightarrow \quad \frac{A_j^{(2)} / \sum_{j'=1}^{P} A_{j'}^{(2)}}{A_j^{(1)} / \sum_{j'=1}^{P} A_{j'}^{(1)}} < \frac{A_k^{(2)} / \sum_{j'=1}^{P} A_{j'}^{(2)}}{A_k^{(1)} / \sum_{j'=1}^{P} A_{j'}^{(1)}} \quad \Leftrightarrow \quad \frac{A_j^{(2)}}{A_j^{(1)}} < \frac{A_k^{(2)}}{A_k^{(1)}}$$

□

*Proof for Proposition 4.* Because fewer than half of all the taxa are differentially abundant, we can be certain that the median absolute abundance fold change is equal to one, namely $\text{Median}\{A_{j'}^{(2)}/A_{j'}^{(1)}\}_{j'=1,2,\cdots,P} = 1$. Based on Proposition 3,

$$R_j^{(2)}/R_j^{(1)} = \text{Median}\{R_{j'}^{(2)}/R_{j'}^{(1)}\}_{j'=1,2,\cdots,P} \quad \Leftrightarrow \quad A_j^{(2)}/A_j^{(1)} = \text{Median}\{A_{j'}^{(2)}/A_{j'}^{(1)}\}_{j'=1,2,\cdots,P}$$

It immediately follows that

$$R_j^{(2)}/R_j^{(1)} = \text{Median}\{R_{j'}^{(2)}/R_{j'}^{(1)}\} \quad \Leftrightarrow \quad A_j^{(2)} = A_j^{(1)}$$

□

### S2 Computational Details of Tobit Model

The Tobit model [10] is the core of ADAPT. When analyzing rare taxa, many observations are left-censored, which may cause the standard estimation method for Tobit models to fail to converge. We adopt several computational heuristics to guarantee the Tobit model estimation's consistency and numerical stability.

#### S2.1 Reparameterization

We use $\{Y_i, \delta_i\}_{i=1,2,\cdots,N}$ to denote the value and censorship indicator of $N$ samples. The censorship indicator $\delta_i$ equals zero if the observation is left censored and one otherwise. Each observation has its corresponding vector of covariates $\boldsymbol{x}_i$. Suppose there are $P$ covariates, including the intercept. The log-likelihood of the Tobit model is

$$\ell = \sum_{i=1}^{N} \delta_i \log \phi \left( \frac{Y_i - \boldsymbol{x}_i \boldsymbol{\beta}}{\sigma} \right) + \sum_{i=1}^{N} (1-\delta_i) \log \Phi \left( \frac{Y_i - \boldsymbol{x}_i \boldsymbol{\beta}}{\sigma} \right) \tag{1}$$

where $\phi(\cdot)$ and $\Phi(\cdot)$ represent the probability density and cumulative distribution of standard normal distribution. This parameterization of the Tobit model with $\boldsymbol{\beta} = \begin{bmatrix} \beta_0 & \beta_1 & \cdots & \beta_{P-1} \end{bmatrix}^\top$ as the effect sizes and $\boldsymbol{\sigma}$ as the scale may lead to multiple solutions during numerical optimization of maximum likelihood estimation [11]. This problem can be avoided if we choose to reparameterize $\boldsymbol{\beta}$ and $\sigma$ with $\boldsymbol{\rho} = \boldsymbol{\beta}/\sigma$ and $\omega = 1/\sigma$ [12]. This parameterization guarantees that the log-likelihood is globally concave and has only one maximum likelihood estimate. The log-likelihood of the Tobit model thus becomes

$$\ell = \sum_{i=1}^{N} \delta_i \log \phi \left( \omega Y_i - \boldsymbol{x}_i \boldsymbol{\rho} \right) + \sum_{i=1}^{N} (1-\delta_i) \log \Phi \left( \omega Y_i - \boldsymbol{x}_i \boldsymbol{\rho} \right) \tag{2}$$

The hypothesis test for one effect size in $\boldsymbol{\beta}$ such as $H_0 : \beta_1 = 0$ against $H_1 : \beta_1 \neq 0$ is equivalent to $H_0 : \rho_1 = 0$ against $H_1 : \rho_1 \neq 0$. The MLE of $\boldsymbol{\beta}$ is $\hat{\boldsymbol{\beta}} = \hat{\boldsymbol{\rho}}/\hat{\omega}$.

#### S2.2 Fisher's Score and Information for log likelihood

We introduce the notation $Z_i = \omega Y_i - \boldsymbol{x}_i^\top \boldsymbol{\rho}$ to simplify the log-likelihood in formula 2 to

$$\ell = \sum_{i=1}^{N} \delta_i \log \phi \left( Z_i \right) + \sum_{i=1}^{N} (1-\delta_i) \log \Phi \left( Z_i \right) \tag{3}$$

The partial derivatives of $Z_i$ over $\omega$ and $\boldsymbol{\rho}$ are

$$\partial Z_i / \partial \omega = Y_i \qquad \partial Z_i / \partial \boldsymbol{\rho} = -\boldsymbol{x}_i \tag{4}$$

The Fisher's score and information of log-likelihood contains partial derivatives of $\ell$ over $Z_i$

$$\frac{\partial \ell}{\partial Z_i} = -\delta_i Z_i + \frac{1-\delta_i}{\sqrt{2\pi}} \cdot \Phi^{-1}(Z_i) \cdot \exp(-\frac{1}{2} Z_i^2) \tag{5}$$

$$\frac{\partial^2 \ell}{\partial Z_i^2} = -\delta_i - \frac{1-\delta_i}{2\pi} \cdot \Phi^{-2}(Z_i) \cdot \exp(-Z_i^2) - \frac{1-\delta_i}{\sqrt{2\pi}} \cdot Z_i \cdot \Phi^{-1}(Z_i) \cdot \exp(-\frac{1}{2} Z_i^2) \tag{6}$$

$$\frac{\partial^3 \ell}{\partial Z_i^3} = \frac{1-\delta_i}{\sqrt{2} \cdot \pi^{3/2}} \cdot \Phi^{-3}(Z_i) \cdot \exp(-\frac{3}{2} Z_i^2) + \frac{3(1-\delta_i)}{2\pi} Z_i \cdot \Phi^{-2}(Z_i) \cdot \exp(-Z_i^2) -$$

$$\frac{1-\delta_i}{\sqrt{2\pi}} \cdot \Phi^{-1}(Z_i) \cdot \exp(-\frac{1}{2}Z_i^2) + \frac{1-\delta_i}{\sqrt{2\pi}} \cdot Z_i^2 \cdot \Phi^{-1}(Z_i) \cdot \exp(-\frac{1}{2}Z_i^2) \tag{7}$$

Fisher's score is the first derivative of log-likelihood $\ell$ over each parameter of $\boldsymbol{\rho}$ and $\omega$. Denote the score vector in the log likelihood as $\boldsymbol{U} = \begin{bmatrix} \boldsymbol{U}_{\boldsymbol{\rho}}^\top & U_\omega \end{bmatrix}^\top$. The first-order partial derivatives are

$$\boldsymbol{U}_{\boldsymbol{\rho}} = \frac{\partial \ell}{\partial \boldsymbol{\rho}} = \sum_{i=1}^{N} \frac{\partial \ell}{\partial Z_i} \cdot \frac{\partial Z_i}{\partial \boldsymbol{\rho}} = -\sum_{i=1}^{N} \frac{\partial \ell}{\partial Z_i} \boldsymbol{x}_i \tag{8}$$

$$U_\omega = \sum_{i=1}^{N} \frac{\delta_i}{\omega} + \sum_{i=1}^{N} \frac{\partial \ell}{\partial Z_i} \cdot \frac{\partial Z_i}{\partial \omega} = \sum_{i=1}^{N} \frac{\delta_i}{\omega} + \sum_{i=1}^{N} \frac{\partial \ell}{\partial Z_i} \cdot Y_i \tag{9}$$

The information matrix $\boldsymbol{I}$ equals the negative hessian matrix $\boldsymbol{H}$. The Hessian matrix $\boldsymbol{H}$ represents the second order partial derivatives of log-likelihood $\ell$ over all the parameters in $\boldsymbol{\rho}$ and $\omega$

$$\boldsymbol{I} = -\boldsymbol{H} = \begin{bmatrix} -\frac{\partial^2 \ell}{\partial \boldsymbol{\rho} \partial \boldsymbol{\rho}^\top} & -\frac{\partial^2 \ell}{\partial \boldsymbol{\rho} \partial \omega} \\ -\frac{\partial^2 \ell}{\partial \omega \partial \boldsymbol{\rho}^\top} & -\frac{\partial^2 \ell}{\partial \omega^2} \end{bmatrix} \tag{10}$$

The four components in $\boldsymbol{I}$ are

$$\boldsymbol{I}_{\boldsymbol{\rho}\boldsymbol{\rho}} = -\frac{\partial^2 \ell}{\partial \boldsymbol{\rho} \partial \boldsymbol{\rho}^\top} = -\sum_{i=1}^{N} \frac{\partial^2 \ell}{\partial Z_i^2} \cdot \boldsymbol{x}_i \boldsymbol{x}_i^\top \tag{11}$$

$$\boldsymbol{I}_{\omega\boldsymbol{\rho}}^\top = \boldsymbol{I}_{\boldsymbol{\rho}\omega} = -\frac{\partial^2 \ell}{\partial \boldsymbol{\rho} \partial \omega} = \sum_{i=1}^{N} \frac{\partial^2 \ell}{\partial Z_i^2} \cdot Y_i \boldsymbol{x}_i \tag{12}$$

$$I_{\omega\omega} = -\frac{\partial^2 \ell}{\partial \omega^2} = -\sum_{i=1}^{N} \frac{\partial^2 \ell}{\partial Z_i^2} \cdot Y_i^2 + \sum_{i=1}^{N} \frac{\delta_i}{\omega^2} \tag{13}$$

#### S2.3 Firth Penalized likelihood

There are situations where a taxon is not observed in any samples from one specific condition. These situations are called complete separation. The complete separation will lead to monotone log-likelihood, and the Newton-Raphson algorithm for maximum likelihood estimation will not converge. To avoid these numerical issues, we add a Firth penalty [13, 14] to the log-likelihood and find the MLE of the penalized likelihood as our parameter estimate. The Firth penalty $\ell_F$ can be understood as Jeffrey's prior on the parameters. The MLE of the penalized likelihood $\ell^\star$ is almost the same as the MLE for $\ell$ except when the sample size is small and/or the existence of a taxon displays patterns of complete separation.

$$\ell^\star = \ell + \ell_F = \ell + \frac{1}{2} \log |\boldsymbol{I}| \tag{14}$$

The Fisher's score $\boldsymbol{U}^\star = \boldsymbol{U} + \boldsymbol{U}_F$. We have derived $\boldsymbol{U}$ in formula 8 and 9 of the previous section. The partial derivative of $\ell_F$ for $\rho_j$ $(j = 0, 1, 2, \cdots, P-1)$ and $\omega$ is

$$U_{F,\rho_q} = \frac{\partial \ell_F}{\partial \rho_q} = \frac{1}{2} \mathrm{Tr} \left( \boldsymbol{I}^{-1} \frac{\partial \boldsymbol{I}}{\partial \rho_q} \right) \tag{15}$$

$$U_{F,\omega} = \frac{\partial \ell_F}{\partial \omega} = \frac{1}{2} \mathrm{Tr} \left( \boldsymbol{I}^{-1} \frac{\partial \boldsymbol{I}}{\partial \omega} \right) \tag{16}$$

The partial derivative of information matrix $\boldsymbol{I}$ over $\rho_j$ and $\omega$ is derived based on the four components in formula 11, 12 and 13. In terms of $\partial \boldsymbol{I}/\partial \rho_q$,

$$\frac{\partial \boldsymbol{I}}{\partial \rho_q} = \begin{bmatrix} \frac{\partial \boldsymbol{I}_{\boldsymbol{\rho}\boldsymbol{\rho}}}{\partial \rho_q} & \frac{\partial \boldsymbol{I}_{\boldsymbol{\rho}\omega}}{\partial \rho_q} \\ \frac{\partial \boldsymbol{I}_{\omega\boldsymbol{\rho}}^{\top}}{\partial \rho_q} & \frac{\partial \boldsymbol{I}_{\omega\omega}}{\partial \rho_q} \end{bmatrix} \tag{17}$$

$$\frac{\partial \boldsymbol{I}_{\boldsymbol{\rho}\boldsymbol{\rho}}}{\partial \rho_q} = \sum_{i=1}^{N} \frac{\partial^3 \ell}{\partial Z_i^3} x_{iq} \cdot \boldsymbol{x}_i \boldsymbol{x}_i^{\top} \qquad \frac{\partial \boldsymbol{I}_{\boldsymbol{\rho}\omega}}{\partial \rho_q} = -\sum_{i=1}^{N} \frac{\partial^3 \ell}{\partial Z_i^3} x_{iq} Y_i \cdot \boldsymbol{x}_i \qquad \frac{\partial I_{\omega\omega}}{\partial \rho_q} = \sum_{i=1}^{N} \frac{\partial^3 \ell}{\partial Z_i^3} x_{iq} Y_i^2 \tag{18}$$

In terms of $\partial \boldsymbol{I}/\partial \omega$,

$$\frac{\partial \boldsymbol{I}}{\partial \omega} = \begin{bmatrix} \frac{\partial \boldsymbol{I}_{\boldsymbol{\rho}\boldsymbol{\rho}}}{\partial \omega} & \frac{\partial \boldsymbol{I}_{\boldsymbol{\rho}\omega}}{\partial \omega} \\ \frac{\partial \boldsymbol{I}_{\omega\boldsymbol{\rho}}^{\top}}{\partial \omega} & \frac{\partial \boldsymbol{I}_{\omega\omega}}{\partial \omega} \end{bmatrix} \tag{19}$$

$$\frac{\partial \boldsymbol{I}_{\boldsymbol{\rho}\boldsymbol{\rho}}}{\partial \omega} = -\sum_{i=1}^{N} \frac{\partial^3 \ell}{\partial Z_i^3} Y_i \cdot \boldsymbol{x}_i \boldsymbol{x}_i^{\top} \qquad \frac{\partial \boldsymbol{I}_{\boldsymbol{\rho}\omega}}{\partial \omega} = \sum_{i=1}^{N} \frac{\partial^3 \ell}{\partial Z_i^3} Y_i^2 \cdot \boldsymbol{x}_i \qquad \frac{\partial I_{\omega\omega}}{\partial \omega} = -\sum_{i=1}^{N} \frac{\partial^3 \ell}{\partial Z_i^3} Y_i^3 - \sum_{i=1}^{N} \frac{2\delta_i}{\omega^3} \tag{20}$$

#### S2.4 BFGS Algorithm

We plan to use the Newton-Raphson algorithm to calculate the MLE of the penalized log-likelihood, but the analytic form of the second-order derivative is hard to derive. Therefore, we resort to the BFGS algorithm [15], a quasi-Newton method. We initialize all entries in $\boldsymbol{\rho}$ to be zero except for the intercept. The intercept $\rho_0$ has initial value $\bar{\boldsymbol{Y}}/\mathrm{SD}(\boldsymbol{Y})$. The inverse scale $\omega$ has initial value $1/\mathrm{SD}(\boldsymbol{Y})$. We set up the initial second-order derivative to be equal to the Hessian matrix of the standard log-likelihood of the Tobit model. The initialization approximates the second-order derivative of the penalized likelihood. The update of the Hessian matrix during each quasi-Newton iteration relies on the approximate Hessian in the previous step, the most recent step size of the parameters, and the most recent change of the first derivatives. The details of the BFGS algorithm are depicted in pseudocode 1

**Algorithm 1** BFGS algorithm for estimating the Tobit model parameters

**Input** Response values of $N$ samples $\boldsymbol{Y}$ , Covariate matrix $\boldsymbol{X}$ with dimension $N \times P$, Taxon existence indicator vector $\boldsymbol{\delta}$ of length $N$, Convergence tolerance threshold $\epsilon$

```
r ← 0                                                         ▷ Iteration counter
ρ^(r) ← 0                                                     ▷ Initialize ρ
ρ_0^(r) ← Ȳ/SD(Y)                                              ▷ Initialize intercept
ω^(r) ← 1/SD(Y)                                               ▷ Initialize inverse scale
θ^(r)⊤ ← [ρ^(r)⊤  ω^(r)]                                      ▷ Concatenate all the parameters
Calculate ℓ*^(r) based on formula 2, 10, 14 and their dependents with θ^(r)    ▷ Initial log likelihood
Calculate U*^(r) based on formula 8, 9, 15, 16 and their dependents with θ^(r) ▷ Initial Fisher score
Calculate B^(r) based on formula 10 and its dependents        ▷ Initialize approximate hessian matrix
do
    r ← r + 1
    Δθ ← −B^(r−1)^−1 U^(r−1)
    θ^(r) ← θ^(r−1) + Δθ                                      ▷ Quasi Newton Update
    Calculate ℓ*^(r) based on formula 2, 10, 14 and their dependents with θ^(r)
    Calculate U*^(r) based on formula 8, 9, 15, 16 and their dependents with θ^(r)
    ΔU ← U*^(r) − U*^(r−1)
    B^(r) ← B^(r−1) + (ΔU ΔU⊤)/(ΔU⊤ Δθ) − (B^(r−1) Δθ Δθ⊤ B^(r−1))/(Δθ⊤ B^(r−1) Δθ)   ▷ BFGS update of the approximate hessian
    Δℓ ← ℓ*^(r) − ℓ*^(r−1)
while |Δℓ/ℓ*^(r−1)| > ε
```

**Output** Estimated parameters $\boldsymbol{\rho}^{(r)}$, $\omega^{(r)}$ and penalized log likelihood $\ell^{\star(r)}$
